# ACE2 fragment as a decoy for novel SARS-Cov-2 virus

**DOI:** 10.1101/2020.04.06.028647

**Authors:** Fabiana Renzi, Dario Ghersi

## Abstract

Novel SARS-Cov-2 enters human cells via interaction between the surface spike (S) glycoprotein and the cellular membrane receptor angiotensin-converting enzyme 2 (ACE2). Using a combination of comparative structural analyses of the binding surface of the S protein to ACE2, docking experiments, and molecular dynamics simulations we computationally identified a minimal, stable fragment of ACE2. This fragment binds to the S protein, is soluble, and appears not to bind to the physiological ligand angiotensinII. These results suggest a possible use of the ACE2 fragment as a decoy that could interfere with viral binding by competition.

## Introduction

The entry of the novel SARS-Cov-2 into the host cell is mediated by the interaction between the viral transmembrane spike (S) glycoprotein and the cellular membrane receptor angiotensin-converting enzyme 2 (ACE2) (1). The S glycoprotein is synthesized as a precursor of about 1,300 amino acids that is then cleaved into an amino N-terminal S1 subunit of about 700 amino acids and a carboxyl C-terminal S2 subunit of about 600 amino acids. Three S1-S2 heterodimers assemble to form a trimer protruding from the viral surface. The S1 subunit contains a receptor-binding domain (RBD), while the S2 subunit contains a hydrophobic fusion peptide. Upon receptor binding, the S1 subunit is cleaved, and the fusion S2 subunit undergoes a conformational rearrangement to form a six-helix bundle that mediates viral and cellular membrane fusion (2).

The angiotensin-converting enzyme (ACE)-related carboxypeptidase, ACE2, is a type I integral membrane protein of about 805 amino acids that contains one HEXXH + E zinc-binding consensus sequence. ACE2 is a close homolog of the somatic angiotensin-converting enzyme (ACE; EC 3.4.15.1), a peptidyl dipeptidase that plays an important role in the renin-angiotensin system. ACE2 sequence includes an N-terminal signal sequence (amino acids 1 to 18), a potential transmembrane domain (amino acids 740 to 763), and a potential metalloprotease zinc-binding site (amino acids 374 to 378, HEMGH). The internal cavity hosts the angiotensinI substrate (consisting of aminoacids 1 to 10 of the angiotensinogen precursor) that ACE2 converts into angiotensinII (amino acids 1 to 8) (3, 4). ACE2 is expressed mainly in heart, kidney, testis, smooth muscle, and in coronary vessels and it seems to increase in lately differentiated epithelial tissues (5). The expression of ACE2 seems inversely regulated by the expression levels of ACE, a key regulator of blood pressure and the target of the pharmacological ACE inhibitors that control blood pressure (6).

X-ray and cryo-EM structures of complexes between the viral S protein from different Coronaviruses both alone and in complex with ACE2 have been solved (7–11). Here we analyzed the crystallographic and cryo-Electron Microscopy (cryo-EM) structures of the S-proteins from SARS-Cov and SARS-Cov-2, human ACE2, and their complexes. We studied the structural features of the binding of ACE2 to S-protein, and performed molecular docking experiments in order to determine the interaction energy between the S protein and ACE2 residues.

We identified a minimal ACE2 fragment consisting of two *α*-helices that retains most of the key interactions with the S protein without interfering with its physiological ligand angiotensinII. Using molecular dynamics (MD) simulations we confirmed that the peptide remains stable in complex with the S-protein. Further, the peptide appears to be stable in an aqueous environment. We conclude by highlighting the potential therapeutic applications of this peptide in terms of scalability, lack of toxicity and immunogenicity, possible administration routes, and potential applications in diagnostic tests.

## Results

### Structural determinants of ACE2 receptor binding to the SARS-Cov-2 and SARS-Cov S-protein

Using the coordinates of the recently solved structure of the complex (`pdb:6VW1`), we determined the per-residue interaction energies between the human ACE2 protein and the SARS-Cov-2 S-protein to ACE2 with `RosettaDock` (12, 13). The results are shown in Figure 1A and 1B, where we highlight the key residues involved in binding. Table 1 lists the interacting amino acids between the human receptor ACE2 and viral proteins from SARS-Cov-2 and SARS-Cov. In addition, we also report the interaction energy values between SARS-Cov-2 and ACE2 computed with `RosettaDock`.

**Table 1.** Interaction energy breakdown for ACE2 residues in complex with SARS-Cov2. The interaction energy values were computed with `RosettaDock`. ACE2 residues are color-coded as in Figure 2, with red for residues with an interaction energy < -1 and yellow for residues with an interaction energy between 0 and -1. The corresponding residues in SARS-Cov (as determined by structural superposition with SARS-Cov-2) are included as well.

| Human ACE2 | Interaction energy | SARS-Cov-2 S-prot. | SARS-Cov S-prot. |
| --- | --- | --- | --- |
| **Alfa helix 1 (-11.3 Rosetta Energy Units)** |  |  |  |
| Ser 19 | -0.018 | Gln 474 | Ser 461 |
| Ser 19 | -0.446 | Ala 475 | Pro 462 |
| Ser 19 | -0.215 | Gly 476 | • |
| Gln24 | -0.602 | Ala 475 | Pro 462 |
| Gln24 | -1.143 | Gly 476 | • |
| Gln24 | -0.085 | Phe 486 | Leu 472 |
| Gln24 | -0.3 | Asn 487 | Asn 473 |
| Gln24 | -0.194 | Tyr 489 | Tyr 475 |
| Thr 27 | -0.78 | Tyr 456 | Tyr 475 |
| Thr 27 | -0.392 | Tyr 473 | Phe 460 |
| Thr 27 | -0.896 | Ala 475 | Pro 462 |
| Thr 27 | -0.95 | Tyr 489 | Tyr 475 |
| Phe 28 | -0.028 | Phe 486 | Leu 472 |
| Phe 28 | -0.069 | Asn 487 | Asn 473 |
| Phe 28 | -0.279 | Tyr 489 | Tyr 475 |
| Asp 30 | -0.05 | Leu 455 | Tyr 442 |
| Lys 31 | -0.445 | Leu 455 | Tyr 442 |
| Lys 31 | -0.333 | Phe 456 | Leu 443 |
| Lys 31 | -0.013 | Glu 484 | Thr 468 |
| Lys 31 | -1.365 | Tyr 489 | Tyr 475 |
| Lys 31 | -0.416 | Gln 493 | Gln 479 |
| His 34 | -0.155 | Val 417 | Val 404 |
| His 34 | -1.766 | Leu 455 | Tyr 442 |
| His 34 | -0.065 | Gln 493 | Gln 479 |
| Glu 35 | -0.039 | Gln 493 | Gln 479 |
| Asp 38 | -0.088 | Gly 496 | Gly 482 |
| Tyr 41 | -0.010 | Gln 498 | Tyr 484 |
| Tyr 41 | -0.111 | Asn 501 | Thr 487 |
| Tyr 41 | -0.031 | Gly 502 | Gly 488 |
| Leu 45 | -0.036 | Thr 500 | Thr 486 |
| **Alfa helix2 (-8.6 Rosetta Energy Units)** |  |  |  |
| Leu 79 | -1.225 | Phe 486 | Leu 472 |
| Ala 80 | -0.035 | Phe 486 | Leu 472 |
| Met 82 | -4.374 | Phe 486 | Leu 472 |
| Tyr 83 | -1.579 | Phe 486 | Leu 472 |
| Tyr 83 | -1.383 | Asn 487 | Asn 473 |
| **Small downstream alfa helix** |  |  |  |
| Asn 330 | -0.032 | Thr 500 | Thr 486 |
| **Beta hairpin (-2.9 Rosetta Energy Units)** |  |  |  |
| Lys 353 | -0.023 | Gln 498 | Tyr 484 |
| Lys 353 | -0.044 | Asn 501 | Thr 487 |
| Lys 353 | -0.195 | Gly 502 | Gly 488 |
| Lys 353 | -2.539 | Tyr 505 | Tyr 491 |
| Gly 354 | -0.053 | Tyr 505 | Tyr 491 |

**Fig. 1.**
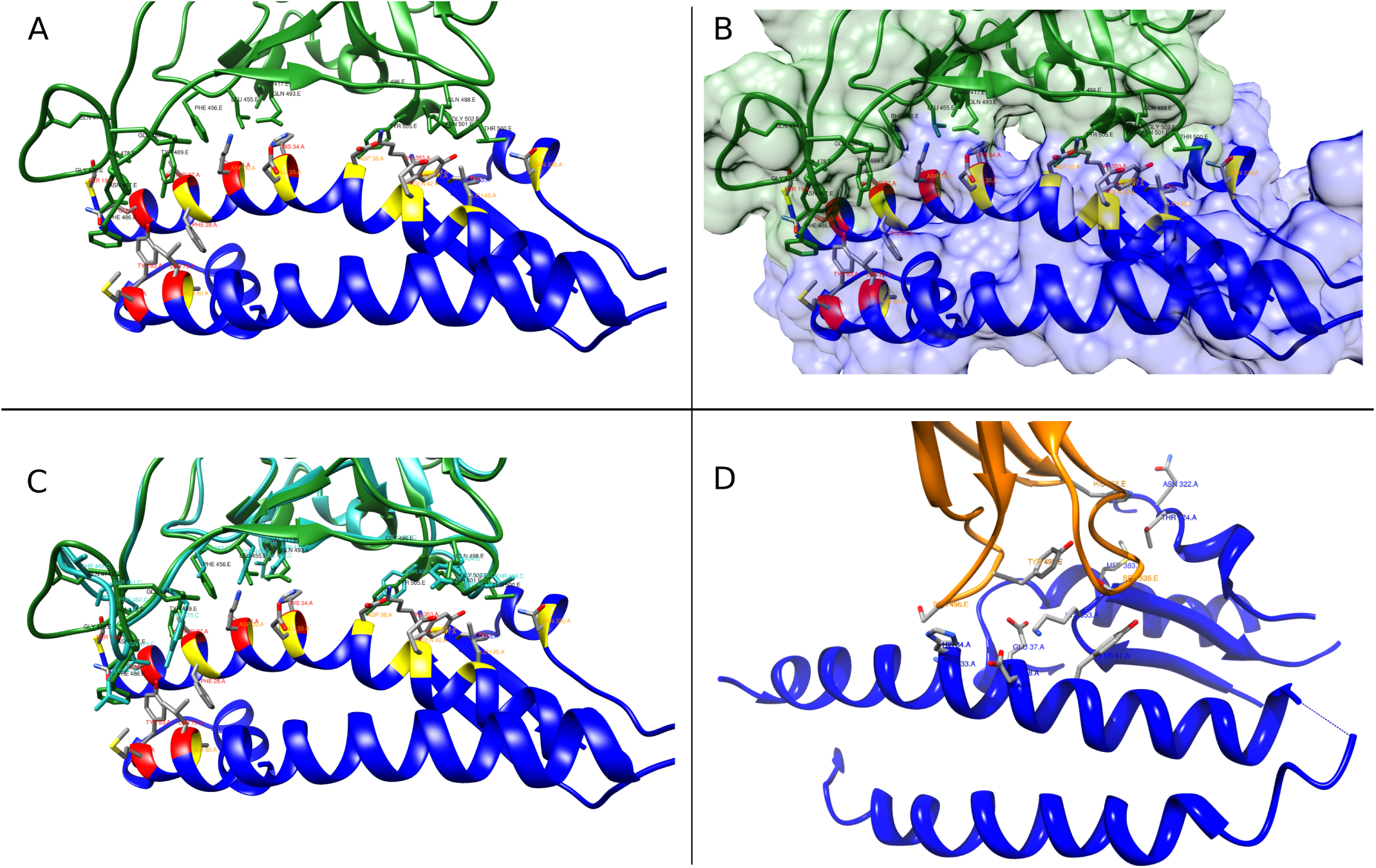
Structural analysis of viral S-proteins binding to ACE2. Panels A and B highlight the ACE2 residues that are involved in binding to the SARS-Cov-2 S-protein. The residues are colored by interaction energy as predicted by `RosettaDock` (red: interaction energy < -1 Rosetta Energy Units (REU); yellow: -1 REU < interaction energy < 0). Panel C shows a superposition of SARS-Cov and SARS-Cov-2 in complex with ACE2, while Panel D shows NL63-Cov S-protein in complex with ACE2.

We also compared complexes of ACE2 bound to the C terminal RBD of viral S-protein from SARS-Cov-2 (`pdb:6VW1`) and SARS-Cov (`pdb:2AJF`) using structural superposition, since these complexes are structurally very similar (Figure 1C). To rule out crystal packing artifacts, we also super-imposed the cryo-EM structure of the SARS-Cov complex (`pdb:6ACK`) onto the crystal structures. We used the coordinates of the S-protein in the pre-fusion conformation (conformation 3 with CTD1 open at 111.6°, which is the physiological state when the S protein binds to the ACE2 receptor). The comparison of the crystal and EM structures did not show evident deviations in the tertiary or secondary structure in regions corresponding to the S-protein binding site (not shown).

In addition, we compared the complex of another coronavirus, Cov-NL63 (`pdb:3KBH`), which targets the ACE2 receptor as well. The structure of Cov-NL63 shows limited structural similarity in binding to ACE2 as compared to other SARS viruses (Fig.1D). Cov-NL63 appears to bind to a narrower area of ACE2, centered around Lys353, which interacts with the conserved viral Tyr498.

The amino acids of the SARS-Cov-2 S-protein involved in binding to ACE2 span a poorly structured region from Leu455 to Tyr505, which correspond to the Tyr442-Tyr491 region in SARS-Cov.

The amino acids in the human ACE2 receptor involved in viral protein binding span two back-to-back helices, *α*-helices 1 and 2, from Ser19 to Tyr83. Based on the per-residue energy breakdown obtained with `RosettaDock`, this area contributes almost 90 percent to the total interaction energy. An additional, point-wise binding feature is represented by Lys353 located in the connecting loop of a downstream beta-hairpin. Lys353 is anchored to the N-terminal ACE2 helix by Tyr41 and Asp38, and it contributes almost 10 percent of the total interaction energy. An additional small downstream helix contributes negligibly to the total binding energy.

In addition, we compared these structural binding determinants with the results of mutagenesis experiments performed by Wong et al. on a SARS-Cov fragment of 193 aminoacids, corresponding to residues 318-510 of the full S-protein (14). Of the two loops encompassing aa 452-454 and aa 434-437 in SARS-Cov-2 (corresponding to aa 460-469 and 447-450 in SARS-Cov and shown to be important in the binding to ACE2), only the first comes into contact with Lys31. The second loop is quite far away from the binding surface (not shown), and its involvement in binding might be indirect, probably affecting the folding of the viral protein fragment.

### The ACE2 fragment remains stable in complex with the viral S-protein and is stable in an aqueous environment

In addition to the interaction energy calculations, the comparative structural analyses discussed above suggest that most ACE2 key residues involved in S-protein binding are found on the N-terminal back-to-back *α*-helices 1 and 2. To determine whether the two helices do in fact remain stable in complex with the S-protein, we performed a 100 ns MD simulation of this complex. We visually inspected the trajectory and computed the root mean square deviation (RMSD) over time with respect to the energy-minimized initial conformer (Figure 2A, green curve). Based on these results, we concluded that the double-helix in complex with the S-protein is stable.

**Fig. 2.**
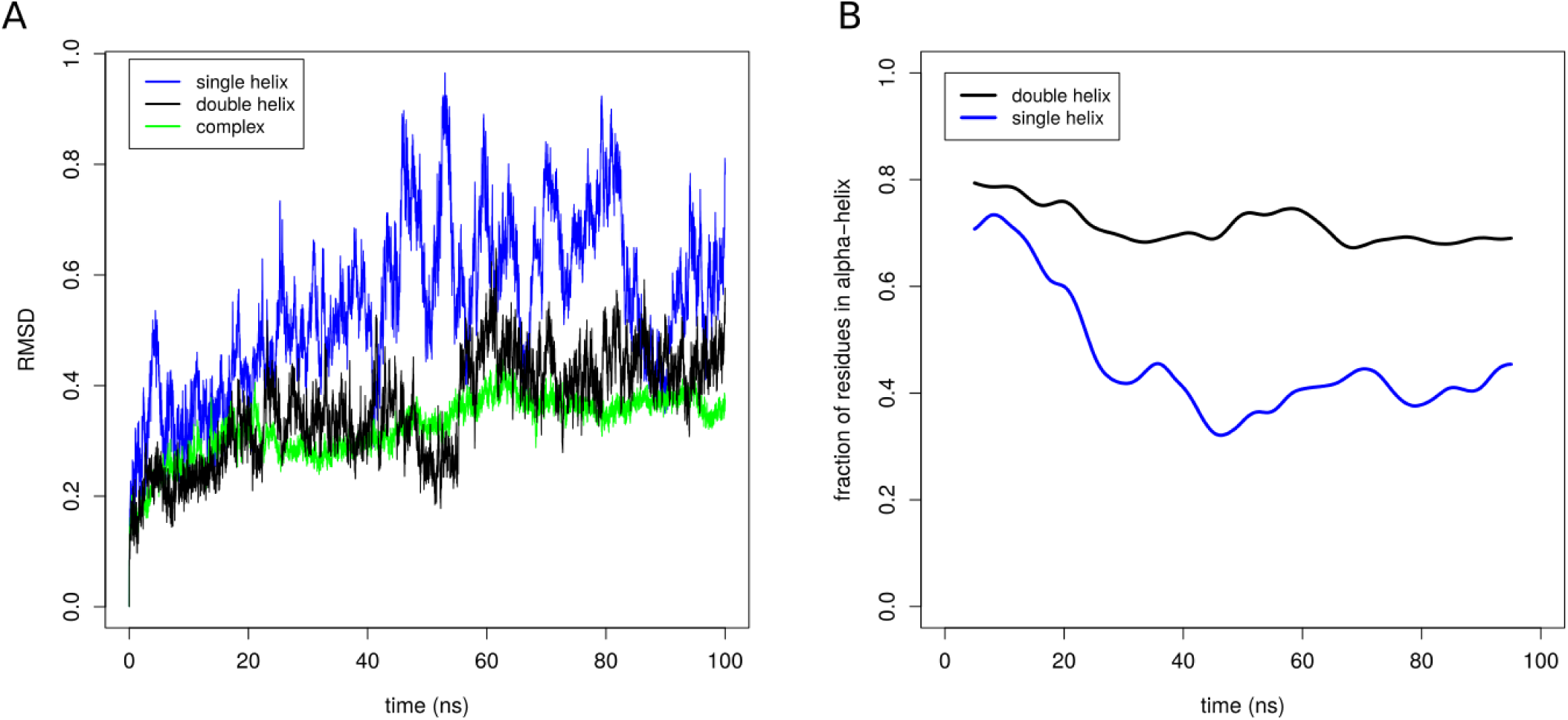
Molecular Dynamics simulations of the ACE2 fragments (single and double helix) and the complex between ACE2 double helix with viral S-protein. Panel A shows the root mean square deviation (RMSD) over time with respect to the initial minimized and equilibrated structures. Panel B shows the fraction of residues in *α*-helix (as determined by DSSP) for the two ACE2 fragments over time.

Further, we determined whether the double-helix is stable in an aqueous environment in the absence of the S-protein, a feature that would be required for a viable inhibitor. We carried out a 100 ns MD simulation of the double-helix alone, visually inspected the trajectory, and compared the RMSD with respect to the initial conformer (Figure 2A, black curve). In addition, we also computed the fraction of residues that remain in *α*-helix conformation over time (Figure 2B, black curve). The results of this analysis indicate that the double helix is stable in an aqueous environment.

We also compared the stability of the double-helix against that of *α*-helix 1. As shown in Figures 2A and 2B (blue curves), the single helix is substantially less stable than the double-helix, with almost half of the residues losing the *α*-helical conformation over time.

### ACE2 fragment does not appear to interfere with the physiological ligand angiotensinII

To determine whether the double-helix fragment has the potential to bind to angion-tensinII, possibly interfering with that physiological system, we performed docking of angiotensinII to ACE2 (Figure 3). As shown by the docking results, the angiotensinII binding site is located in an internal cavity of the receptor, quite far from the surface where the viral S-protein binds. Most of the residues of ACE2 involved in binding to angiotensinII are internal to the core of the protein. Only two amino acids (Asp51 and Ser47) in *α*-helix 1 have appreciable interactions with angiotensinII, and contribute about 15 percent of the total interaction energy as predicted by `RosettaDock`. These results suggest that the double-helix fragment is highly unlikely to interfere with angiotensinII in a physiological context.

**Fig. 3.**
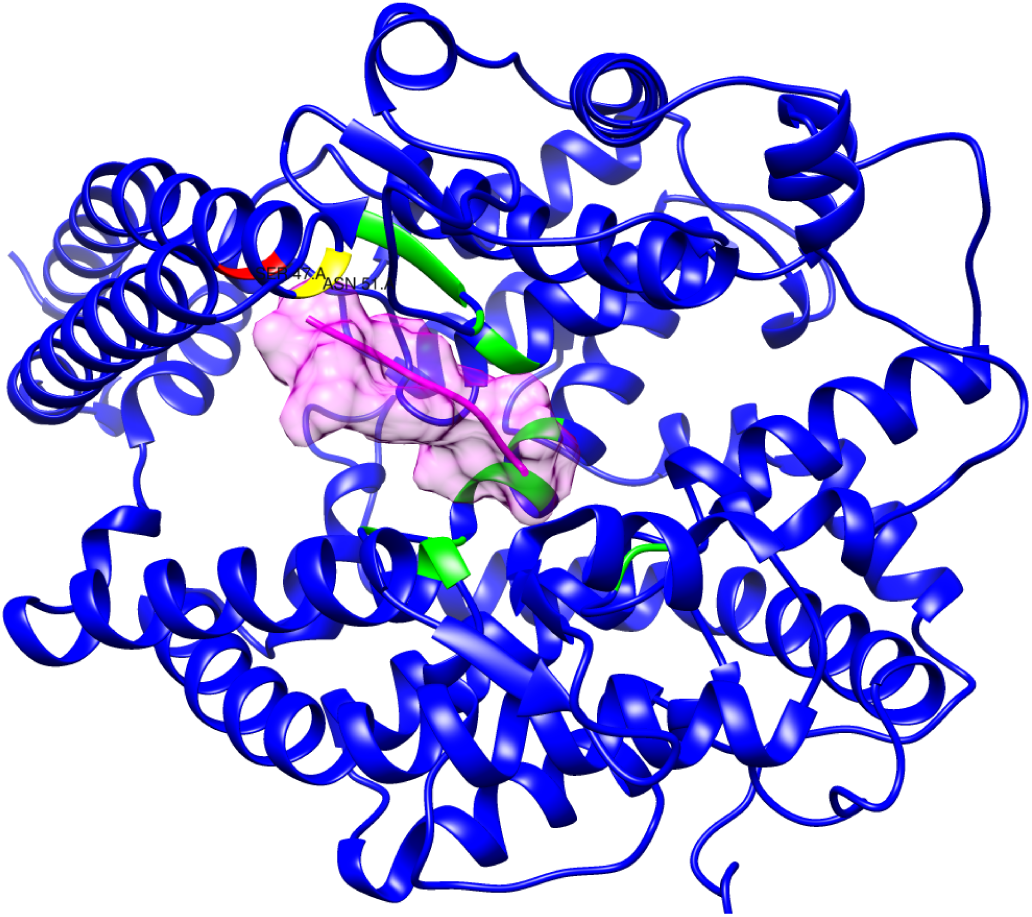
Molecular docking experiment with angiotensinII. The molecule surfaced in pink is angiotensinII docked to ACE2. The residues in red and yellow are the only two positions in the double helix subject of this study that make contact with angiotensinII. The residues highlighted in green represent the bulk of the binding site and lie outside of the double helix fragment.

## Discussion and Conclusions

In the context of a pandemic, when the scientific community and the pharmaceutical industry are hard pressed to discover and develop life-saving treatments in a short time, this work suggest an easy-to-test and potentially viable strategy to interfere with the first step of SARS-Cov-2 and SARS-Cov entry into human cells.

By performing comparative structural analyses, molecular docking, and molecular dynamics simulations we identified the key residues that are involved in the binding between the viral S-protein and the human ACE2 receptor. These key residues are found in the two N-terminal back-to-back helices 1 and 2, spanning about 60 amino acids and contributing most of the predicted interaction energy between ACE2 and the SARS-Cov-2 and SARS-Cov proteins.

An ideal fragment would also include the downstream beta-hairpin carrying the conserved Lys353. However, this residue is too far in the primary structure from the double helices. A cut and seal between these two binding regions is unlikely to work due to the unpredictable folding of the resulting chimeric structure. Therefore, we propose to use the double-helix fragment as a candidate for effectively inhibiting the S-protein.

We also showed that this fragment remains stable in an aqueous environment (as opposed to a shorter fragment comprising only helix 1). Further, the fragment does not appear to interfere with the physiological ligand of ACE2 (angiotensinII), which is part of the renin-angiotensin system involved in the regulation of blood pressure.

We would expect this fragment to be relatively easy to express and purify at a large-scale level^1^. Since this fragment is derived from a protein that is normally expressed in human tissues, the fragment should be relatively safe to use. The fragment might be administered directly by aereosol, avoiding the systemic route and possible degradation of the peptide.

Additionally, the fragment might also be employed for diagnostic purposes. For example, immobilized on a substrate for a Biacore test, the fragment could serve as a quick binding assay to detect viral particles.

## Materials and Methods

### Structural data for comparative analyses

For the structural analyses we used the following pdb structures:

- `6VW1`: crystal structure of the S protein in complex with ACE2 from SARS-CoV-2 (chain A and chain E) (10)
- `6ACK`: cryo-EM structure of the S protein in complex with ACE2 from SARS-CoV (chain D and amino acids 23-512 of chain C, corresponding to CTD1 of S protein; extracted from complex between S-trimer and ACE2) (9)
- `3KBH`: crystal structure of the S protein in complex with ACE2 from CoV-NL63 (chain A and E) (11)
- `4APH`: crystal structure of ACE1 in complex with angiotensinII (15)
- `1R42`: crystal structure of ACE2 (4)

To perform structure superpositions we used the `MatchMaker` function as implemented in `UCSF Chimera` (16).

### Docking experiments

Docking experiments and interaction energy calculations were carried out using `RosettaDock` (12, 13). To obtain the per-residue energy breakdown of the ACE2/S-protein complex we relaxed the structure to relieve clashes using the `flag_input_relax` protocol provided with the `Rosetta` suite. We then optimized the complex by performing 1,000 redocking experiments with the local docking protocol provided in the suite, and selected the pose with the most favorable interaction energy. The per-residue energy breakdown was obtained with the `residue_energy_breakdown` provided in the suite. For the docking experiment with angiontensinII, we used the ligand found in the crystal structure of the complex with the ACE receptor (`pdb:4APH`, chain P) superposed onto the crystal structure of ACE2. After relaxing the complex, we carried out 1000 docking experiments and retained the pose with the most favorable interaction energy.

### Molecular Dynamics Simulations

Molecular dynamics Simulations were performed using GROMACS (17, 18) with the all-atom OPLS force field. Non-protein atoms were removed from the PDB files. The protein structures were placed in a cubic box with 10Å from the box edge and solvated with `spc216.gro`, an equilibrated 3-point solvent model.

Next, we added ions to the system with the `genion` module and performed 50,000 steps of energy minimization. We then carried out equilibration in two steps: NVT (canonical) ensemble followed by NPT (isothermic-isobaric). For the NVT equilibration phase, we set the target temperature to 300 K and inspected the temperature graph over time (100 ps) to make sure the system was stabilized around the target temperature value. For the NPT equilibration phase, we used the Parrinello-Rahman barostat, and inspected the density values over time (100 ps) to check the stability of the system.

After the equilibration steps, we ran a 100 ns production run on a GPU-accelerated machine. All MD simulations were performed with the same protocol and a time step of 2 fs. The trajectories were visually inspected in UCSF Chimera (16), and the RMSD with respect to the initial energy-minimized conformer was obtained with the `rms` module in the GROMACS suite.

We computed the fraction of residues in *α*-helix with an in-house Python program. Secondary structure assignment was obtained with the DSSP program (19).

## ACKNOWLEDGEMENTS

Docking and Molecular Dynamics simulations were performed on a GPU-accelerated high-performance computing machine partly funded by a Nebraska EPSCoR grant to DG. Thanks to Dr Antonello De Santis for informatics support to FR.

## Footnotes

1 Since the inception of this work a manuscript by Zhang et al. (bioRxiv 2020.03.19.999318) has shown that a 23-mer from ACE2 can bind to SARS-Cov-2 S-protein.

